# Quantification of H3.1-nucleosomes using a chemiluminescent immunoassay: a reliable method for neutrophil extracellular trap detection

**DOI:** 10.1101/2025.03.05.641629

**Authors:** Marion Wargnies, Guillaume Rommelaere, Julie Candiracci, Dorian Pamart, Robin Varsebroucq, Florian Jibassia, Finley Serneo, Virginie Laloux, Olivia Thiry, Fanny Lambert, Alison Lobbens, Priscilla Van den Ackerveken, Marielle Herzog

**Author notes:** Corresponding author: (MH).

## Abstract

Neutrophil extracellular traps (NETs) are chromatin-based web-like structures released by activated neutrophils in response to infectious agents. Overproduction or insufficient clearance of NETs contributes to dysfunction of immune response and disease pathogenesis, underlying the importance of early detection and monitoring of NET levels in clinical samples. While existing methods for NETs detection and quantification face limitations, there is a pressing need for a reliable, sensitive, and clinically applicable assay. Since NETs consist of long strains of decondensed chromatin, with nucleosomes as their basic units, we propose circulating H3.1-nucleosomes as biomarkers for NETs detection in clinical plasma samples.

In the initial phase of our study, we confirmed the presence of H3.1-nucleosomes by immunofluorescence and immunoprecipitation experiments in two *in vitro* NET models: neutrophil-like cells differentiated from the HL-60 cell line and primary neutrophils isolated from whole blood, both treated with phorbol 12-myristate 13-acetate to induce NET formation. Subsequently, we developed and analytically validated a chemiluminescent immunoassay for the quantification of circulating H3.1-nucleosomes in plasma. This fully automated assay demonstrates impressive analytical performance in parameters including sensitivity, precision, linearity and reproducibility. Overall, by measuring the H3.1-nucleosome levels in plasma samples from patients suffering from NETs-related diseases compared to healthy donors, we demonstrated the assay’s potential as a groundbreaking diagnostic tool for disease management.

## 1. Introduction

Nucleosomes are the basic structural units of chromatin, consisting of DNA wrapped around a core of histone proteins (H2A, H2B, H3, and H4) (1). In the nucleus, these complexes are involved in chromatin compaction, contributing to gene expression regulation (2, 3). Beyond their intracellular functions, circulating nucleosomes have emerged as significant biomarkers in various pathological conditions due to their association with cellular stress, damage, and death (4-7). Among the different types of cell death, NETosis appears as a particularly intriguing process (8). NETosis is a specialized form of programmed cell death of neutrophils, the most abundant circulating white blood cell and the first line of defense in the innate immune response. NETosis results in the release of neutrophil extracellular traps (NETs), which are composed of long stretches (i.e. 15,000-30,000 bp) of decondensed chromatin decorated with granule proteins such as myeloperoxidase (MPO) and neutrophil elastase (NE) (9, 10). Besides the beneficial role of NETs in the defense against pathogens, excess production or dysregulation of NET release is associated with exacerbated inflammation and organ damage (11-14). The significant contribution of NETs in various pathologies including sepsis, COVID-19, rheumatoid arthritis, systemic lupus erythematosus, and others (15-25) highlights the importance of detecting and monitoring NET levels in clinical samples (26-28). Traditionally, NETs are detected using methods such as fluorescence microscopy or enzyme-linked immunosorbent assays (ELISA) targeting NETs components such as MPO, cell-free DNA (cfDNA), citrullinated histones H3 (H3Cit), and MPO-DNA complexes (29-34). Although these methods have been described in the literature to detect NETs *in vitro* and *in vivo*, they have several limitations in clinical applications, particularly regarding sensitivity, standardization and automation. This highlights the need for a robust and analytically validated method for NET detection in clinical settings (35, 36). Since NETs are composed of decondensed chromatin, and nucleosomes are inherent constituents of this structure, we hypothesize that circulating nucleosomes − particularly those containing the H3.1 histone variant − could serve as biomarkers for NET detection in clinical samples (37-39). Investigating nucleosomes offers several advantages over other NET components, such as free histones or cfDNA, due to their inherent structural stability, which protects them from rapid degradation in the bloodstream (40). Additionally, unlike histone post-translational modifications (PTMs) that may be affected by enzymatic activity or clipping events, nucleosomal detection is less influenced by these variables.

In this study, we first confirmed the presence of H3.1-nucleosomes in NETs using two complementary *in vitro* models; the well-established DMSO-differentiated HL-60 cells (41, 42) and PMA-stimulated primary human neutrophils (43). Immunofluorescence microscopy confirmed the presence of H3.1-nucleosomes in the NETs structures in both models. To further validate these findings, immunoprecipitation (IP) experiments demonstrated the physical association of H3.1-nucleosomes with other NETs components in HL-60-derived NETs. These results collectively support the relevance of H3.1-nucleosome quantification as a biomarker for NETs.

Building on this, we developed an automated and standardized chemiluminescent immunoassay (ChLIA) for the detection of circulating H3.1-nucleosomes in plasma samples. This fully automated assay employs an anti-histone H3.1 antibody in capture on the solid phase, consisting of magnetic particles, in combination with a conformational anti-nucleosome antibody coupled with acridinium ester molecules in detection. Analytical validation demonstrated that the assay is highly sensitive, precise, linear, and reproducible.

Finally, we evaluated the clinical utility of this H3.1-nucleosome immunoassay by measuring circulating H3.1-nucleosome levels in plasma samples from patients suffering from diverse NETs-associated diseases compared with healthy donors. The results showed significantly elevated levels of circulating H3.1-nucleosomes in patients with NETs-related diseases compared with the control population, confirming the assay’s potential for monitoring NET levels in clinical settings. Overall, this study demonstrates the development of an analytically validated quantitative NET assay and highlights the strong potential of the H3.1-nucleosome immunoassay as a valuable diagnostic tool for NETs-associated conditions.

## 2. Materials and Methods

### 2.1. HL-60 cell line culture and NETosis induction

HL-60 cells were purchased from ATCC (CCL240; ATCC; Manassas, USA). The cells were cultured in Iscove’s Modified Dulbecco’s Medium (IMDM) (31980048; ThermoFisher scientific; Waltham, USA) glutaMAX (31980048; Gibco, ThermoFisher scientific, Waltham, USA) supplemented with 15% fetal bovine serum (A4766801; Gibco, ThermoFisher scientific, Waltham, USA) and a penicillin/streptomycin solution (5.000U: 15070063; Gibco, ThermoFisher scientific, Waltham, USA). Cultures were maintained in a humidified atmosphere with 5% CO_2_ at 37°C. To synchronize the cell population prior to differentiation, the percentage of FBS was reduced to 1%, and the cells were incubated in this restricted medium for 48 hours. For differentiation into granulocyte-like cells, HL-60 cells were then incubated with 1.5% dimethyl sulfoxide (DMSO; D2650; Merck Life Sciences BV; Hoeilaart, Belgium) for 5 days. After the differentiation period, cell viability was assessed using the Eve^TM^ plus cell automatic counter (NanoEnTek; Seoul, Korea).

For NETosis induction, differentiated cells were resuspended at a concentration of 10^6^ cells/mL in IMDM without additives. The resuspended cells were then seeded in 6-well plates with 2 mL of cell suspension per well (2 x 10^6^ cells/well). To induce NETosis, 100 nM phorbol 12-myristate 13-acetate (PMA; J63916; ThermoFisher scientific, Waltham, USA) was added to each well, except for control condition, and the cells were incubated for 5 hours. For control conditions, cells were incubated with an equivalent volume of DMSO, the PMA diluent. After the 5-hours incubation period, the cells adhering to the bottom of the wells were scraped, and the entire contents of the wells were collected into Falcon conical tubes (430291; Corning Life Sciences; Corning, USA). The tubes were then centrifuged at 200 g for 2 minutes to remove cell debris. The supernatant, containing NETs extruded in the IMDM, was collected. To increase the stability of the samples during their storage at -80°C, 20% glycerol (G5516-100 mL; Merck Life Sciences BV; Hoeilaart, Belgium) was finally added in the tubes.

### 2.2. Human Neutrophil isolation and NETosis induction

Human neutrophil isolation was performed within one hour of blood collection using the MACSxpress Whole Blood Neutrophil Isolation Kit (130-104-434; Myltenil Biotec, Gaithersburg, USA) for humans as previously published (43). At the end of this isolation step, the isolated neutrophils cell pellet was resuspended in prewarmed RPMI medium at 37°C at a concentration of 1-2 x 10^6^ cells/mL for NETosis induction with 100 nM of PMA.

### 2.3. Immunofluorescence Microscopy

HL-60 cells were incubated in 6-well plates containing one coverslip (631-0150; VWR; Radnor, USA) coated with poly-L-lysine per well. During the 5-hour NETosis induction period, cells adhered to the coverslips. Then, post-stimulation, samples were fixed with 1.3% paraformaldehyde (047317.92; ThermoFisher scientific, Waltham, USA) for 30 minutes at room temperature (RT) and then blocked with 5% donkey serum (#D9663; Merck Life Sciences BV, Hoeilaart, Belgium) for 30 minutes at RT. The primary neutrophil cells were incubated at 37°C with 5% CO_2_ 3 hours, followed by fixation with 1% paraformaldehyde for 10 minutes before the immunostaining characterization. DNA was counterstained with 4’,6-diamidino-2-phénylindole (DAPI; D1306; ThermoFisher scientific, Waltham, USA) and cell membranes with Cell Mask^TM^ Orange (C10045; ThermoFisher scientific, Waltham, USA). The coverslips were incubated with primary antibodies directed against myeloperoxidase (MPO) (ab221847; GR3285980-23; Abcam, Cambridge, UK, dilution 1:100), histone H3.1 (Belgian Volition SRL, Isnes, Belgium, dilution 1:100) or nucleosome (Belgian Volition SRL, Isnes, Belgium, dilution 1:50) for 2 hours at RT, protected from the light. This was followed by incubation with secondary antibodies conjugated with Alexa Fluor fluorochromes (dilution 1:1000) (Goat anti-rabbit IgG AF 647; A-21244; ThermoFisher scientific, Waltham, USA) (Goat anti-mouse IgG AF 488; A-11001; ThermoFisher scientific, Waltham, USA) (Donkey anti-mouse IgG AF 647; A-31571; ThermoFisher scientific, Waltham, USA) (Donkey anti-goat IgG AF 488; A-11055; ThermoFisher scientific, Waltham, USA) for 1 hour at RT, protected from the light. After staining, slides were mounted by adding one drop of Fluoromount-G mounting solution (00-4958-02; ThermoFisher scientific; Waltham, USA) on a glass slide and carefully placing a coverslip facing downwards on the medium. Mounted slides were dried overnight at RT, in the dark. NETs derived from differentiated HL-60 cells were visualized using a Zeiss LSM900 Airyscan confocal laser scanning microscope (Zeiss; Oberkochen, Germany) equipped with 63x and 40x magnification objectives. Images were acquired at the Morph-IM platform (UNamur Technology Platform for Morphology and Imaging). For experiments involving NETs generated by isolated primary neutrophils, imaging and analysis were performed using a Nikon AXR Confocal with NSPARC Super Resolution microscope (Nikon; Tokyo, Japan) at the Nikon Imaging Center at UC San Diego.

### 2.4. NETs immunoprecipitation

MyOne Tosylactivated magnetic beads (65501; ThermoFisher scientific; Waltham, USA) were coated with the following antibodies: an anti-histone H3.1 monoclonal antibody (Belgian Volition SRL, Isnes, Belgium) as previously described (44), an anti-nucleosome monoclonal antibody (Belgian Volition SRL, Isnes, Belgium), and an anti-MPO monoclonal antibody (475915; 3460549; Merck Life Sciences BV; Hoeilaart, Belgium). To immunoprecipitate proteins of interest, 500 µL of sample was incubated with 1 mg of coated beads for 1 hour at RT (for anti-H3.1- and anti-MPO-coated beads) or 37°C (for anti-nucleosome-coated beads) on a roller. Following the incubation, the beads retaining the precipitated proteins were pelleted using a magnetic separation rack and were washed three times on a roller. Then, the products of immunoprecipitation (IP) were assessed either by Western blotting for the protein profile or by Agilent 2100 Bioanalyzer (Agilent Technologies; Santa Clara, USA) for the DNA profile.

### 2.5. dsDNA extraction

Following the NETs IP protocol, DNA from both the supernatant post-IP and the immunoprecipitated material was extracted using QIAamp^®^ DSP Circulating NA kit (61504; Qiagen; Venlo, The Netherlands) according to the manufacturer’s instructions.

### 2.6. dsDNA size distribution

The extracted dsDNA from both the supernatant post-IP and the immunoprecipitated materials were assessed using an Agilent 2100 Bioanalyzer with an Agilent High Sensitivity DNA Kit for fragment sizes of 50-7000 bp (5067; Agilent Technologies; Agilent Technologies; Santa Clara, USA), according to manufacturer’s instructions.

### 2.7. Western blotting

Following the IP protocol, the complex of NETs bounded to magnetic particles was incubated for 5 minutes at 95°C in Laemmli buffer (1610747; Bio-Rad, Hercules, USA) containing 10% of 2-mercaptoethanol (M6250; Merck Life Sciences BV; Hoeilaart, Belgium). The magnetic beads were then pelleted on a magnetic separation rack, and the supernatant containing the denaturated proteins was separated using acrylamide gels. 4-20% TGX precast gels (4561094; Bio-Rad, Hercules, USA) were used for electrophoresis under reducing conditions (Tris/Glycine/SDS; 1610772; Bio-Rad; Hercules, USA).

Proteins were transferred onto polyvinylidene difluoride (PVDF) membranes (1704272; Bio-Rad; Hercules, USA) using a semi-dry transfer system (Bio-Rad; Hercules, USA) for 7 minutes at up to 20V. The PVDF membranes were then blocked using Tris Buffer Saline (TBS) containing 1% casein (1706435; Bio-Rad; Hercules, USA) for 30 minutes to 1 hour before being incubated for 2 hours with anti-MPO (antibody recognizing the two subunits α and β of the protein MPO, 2 µg/mL; 475915; 3460549; Merck Life Sciences BV, Hoeilaart, Belgium), anti-histone H3 (0.5 µg/mL; 91297; 3162002; Active Motif, Carlsbad, USA), or anti-histone H4 antibody (1/1000; ab177840; Abcam; 1014591-13; Cambridge, UK) diluted in TBS containing 1% casein.

After the primary antibody incubation, the membrane was washed three times and then incubated with a detection reagent (VeriBlot antibody for IP Detection Reagent, ab131366; 1035165-4; Abcam, Cambridge, UK) for 1 hour at RT. Finally, after three additional washes, the membrane was incubated with SuperSignal™ West Dura Extended Duration Substrate (34076; ThermoFisher scientific; Waltham, USA) and chemiluminescent signal was acquired using the Fusion-FX6 instrument and software (Vilber, Marne-la-Vallée, France).

To reuse the same membrane for different stainings, they were rinsed three times with TBS containing 0.1% of Tween^®^20 (T2700; Merck Life Sciences BV; Hoeilaart, Belgium) (TBS-T) and subjected to a mild stripping protocol. Briefly, the membrane was incubated with stripping buffer for 10 minutes twice, washed with PBS 1x for 10 minutes twice, and washed with TBS containing 0.1% of Tween^®^20 (TBS-T) for 5 minutes twice before being blocked again. The rest of the protocol remained the same as described previously in this section. The efficiency of stripping was checked by incubating the membrane with chemiluminescent detection reagent just before the second blocking step.

### 2.8. Quantification of nucleosomes with the Nu.Q^®^ ChLIA assays

H3.1- and H3R8Cit-nucleosome levels were measured using the Nu.Q^®^ H3.1 (2001-02-01; Belgian Volition SRL, Isnes, Belgium) and Nu.Q^®^ H3R8Cit ChLIA assays (Belgian Volition SRL, Isnes, Belgium), respectively. Both assays were developed on the IDS-i10 automated chemiluminescence immunoanalyzer system (Immunodiagnostic Systems Ltd; Boldon, UK) with magnetic beads technology. Briefly, 50 μL of samples are mixed with an acridinium ester labeled anti-nucleosome detection antibody and incubated for 30 minutes. The conformational antibody ensures that the measured analyte is the nucleosome itself, distinguishing it from individual histones or circulating DNA. Anti-histone H3.1 or H3R8Cit antibody-coated magnetic beads were then added and incubated for an additional 15 minutes. Following incubation, a washing step was performed to remove unbound components. Trigger solutions were added, and the emitted light was recorded using the luminometer. A standard curve specific for each assay was used for calibration. Sample quantification was performed using a 4-parameter logistic curve fitting. The Nu.Q^®^ H3.1 assay quantifies H3.1-nucleosome concentration from 3 to 1200 ng/mL. Samples exceeding 1200 ng/mL were automatically remeasured following an automated 5-fold dilution by the instrument, extending the quantification range up to 6000 ng/mL. The Nu.Q^®^ H3R8Cit assay quantifies H3R8Cit-nucleosome concentration from 5 to 500 ng/mL.

### 2.9. Analytical performances evaluation of the H3.1-nucleosome ChLIA assay

Experimental designs for evaluating the analytical performances of the H3.1-nucleosome assay for K2EDTA plasma measurements followed the recommendations of the CLSI guidelines. Precision was assessed by measuring five samples with different concentration levels, obtained by spiking human K2EDTA plasma samples with native nucleosomes extracted from human cells. The samples were measured, using three lots of reagents over 7 days, with two runs per day and two measurements per run (3 x 7 x 2 x 2 design). Coefficients of variation percentages (CV%) were calculated based on CLSI-EP05A suggested analysis of variance using the Analyse-it software, allowing evaluation of repeatability, within-lot precision, and within-laboratory precision.

Sensitivity limits were determined based on CLSI guideline EP17-A2 suggestions, using three independent lots of reagents. For the limit of blank (LOB), 60 measurements of diluent were conducted over 5 days, and the LOB was calculated as the average value plus 1.645 standard deviation (SD), representing the 95^th^ percentile of the normal distribution. The limit of detection (LOD) was calculated by measuring 10 human K2EDTA plasma samples, ranging from approximately 0 to 10 ng/mL, in triplicates over 5 days. The pooled standard deviation (pooled SD) for all the measurements was determined and the LOD was calculated as follows: LOD = LOB + 1.645 pooled SD of non-blank samples. Finally, the lower limit of quantification (LOQ) was defined for three lots of reagents using a precision profile, setting 15 CV% as the maximum limit of imprecision allowed for precise quantification.

For the evaluation of the linearity, human K2EDTA plasma samples of different concentrations were mixed in equivalent proportions (per 10% levels) and linear curve fitting with variance-based weighting were applied. The linear ranges were established by comparing measured concentrations of the different sample proportions with linear regression expected concentrations. The linearity range defines the range within which the difference between measurement and linear model predictions is less than 10%.

In spike-recovery experiments, three samples were spiked with two concentrations of recombinant nucleosomes and compared with these samples added with the same volume of diluent. The percentage of recovery was evaluated by normalizing the measured concentration of spiked samples with the expected nucleosome concentration (measured concentration of the control + spiked concentration).

Interferences were evaluated by comparing nucleosome concentrations in plasma samples with and without the addition of the following components: bilirubin (conjugated or unconjugated), human serum albumin, triglycerides, cholesterol, or hemoglobin at different concentrations (representing high, non-normal concentrations). For human anti-mouse antibodies and rheumatoid factor, a spike recovery was conducted in samples with known concentrations of these two heterophilic antibodies to verify the absence of false positive or false negative influence.

Cross-reactivity was tested by measuring single H3.1-histone and H3.3-containing nucleosomes, as well as citrullinated recombinant nucleosomes H3R8Cit and H3R2,8,17Cit.

Carryover was investigated by repeatedly measuring a sample of low and a sample of high nucleosome concentrations to ensure that the order of measurement did not influence the precision or accuracy of the measurements.

### 2.10. Study population

The study included 236 samples from control donors, self-certified as healthy donors from EFS (Etablissement Français du Sang) and 302 samples from patients with diseases associated with NETosis such as COVID-19 (n = 80), sepsis suspicion and confirmed (n = 109), cirrhosis or nonalcoholic steatohepatitis (n = 11), cytomegalovirus infection (n = 23), gonorrhea infection (n = 10), myocardial infarction (n = 38), Alzheimer’s disease (n = 12), hepatitis A virus infection, (n = 6), Lyme infection (n = 6), traumatic brain injury (n = 6), and heart failure (n = 1). Diagnoses were provided by biobank suppliers.

Frozen K2EDTA plasma samples used in this study were sourced from certified biobanks, including INOSpecimens, BioIVT, DxBioSamples, BayBiosciences, and QUALIblood and were received anonymized. All specimens were either collected from consented donors or provided as de-identified remnant samples. Each biobank adheres to stringent ethical policies and implements state-of-the-art procedures to ensure the highest ethical standards in its operations in accordance with the Declaration of Helsinki, ensuring the protection of human subjects. For samples obtained from INOSpecimens BioBank (Clermont Ferrand, France), the biobank holds authorization to commercialize residual samples from medical analyses for scientific research purposes (AC-2018-3151, AC-2022-4975). Samples from the other biobanks were retrospectively collected and made available for research in compliance with applicable national regulations. All enrolled patients signed specific informed consent forms for blood draw and related translational research projects. The protocol/study numbers for all the samples tested are as follows: BioIVT (Frankfurt, Germany) 2020-002/1282289; DxBioSamples LLC (San Diego, California, USA) PG-ONC 2003/1; BayBiosciences LLC (Brookline, Massachusetts, USA) BBS-TPP-817-22; QUALIblood (Liège, Belgium) B03920096633 49/2009.

The study is a research study conducted on retrospective biobanked samples. It is not a clinical trial and does not involve the recruitment of new participants. Therefore, ethical approval is not required for this study.

### 2.11. Statistical analysis

Analytical data were analyzed using Analyse-it for Microsoft Excel (version 5.9) software. The linearity of the model was assessed through simple linear regression using linear fit model, allowing ±10% of non-linearity.

Descriptive statistical analysis of the clinical dataset was performed using GraphPad Prism (version 9.5.0, San Diego, CA, USA). The data were subjected to the Kolmogorov– Smirnov normality test. Group comparisons were conducted using Welch’s test and the Mann–Whitney U test for non-parametric data. Specificity, sensitivity, negative predictive value, and positive predictive value were calculated. The area under the curve (AUC) were calculated from the receiver operator characteristic (ROC) curve. The Youden Index (*J*), defined as *J* = *Sensitivity* + *Specificity* − 1, was employed to identify the optimal threshold. Significance values are represented as follows: *, p-value (p) < 0.05; **, p < 0.01; ***, p < 0.001; ****, p < 0.0001.

## 3. Results

### 3.1. The molecular characterization of NETs produced *in vitro* confirms the presence of H3.1-nucleosomes

HL-60 promyeloblastic cells can be differentiated into neutrophil-like cells through exposure to dimethyl sulfoxide (DMSO). Once differentiated, subsequent treatment with phorbol 12-myristate 13-acetate (PMA) induces NETosis. Similarly, PMA triggers NETosis in primary human neutrophils isolated from whole blood. To confirm NETosis induction by PMA treatment, we compared the presence of NET structures in PMA-treated DMSO-differentiated HL-60 cells (DMSO-dHL-60+PMA) with those in untreated cells (DMSO-dHL-60) (S1 Figure). In DMSO-dHL-60 cells, the fluorescent microscopy images showed intact cell membranes stained with Cell Mask^TM^ Orange and intact nuclei marked with DAPI, indicating the absence − or a weak induction − of NETosis. In contrast, PMA-treated cells exhibited cell membrane debris and large extruded DNA filaments, confirming NETosis activation. Then, we analyzed, the putative spatial relationship between NETs and H3.1-nucleosomes in the two *in vitro* NETs models – the DMSO-dHL-60 +PMA model and the Primary Neutrophils +PMA model –, using antibodies directed against the protein MPO, the histone variant H3.1, the citrullinated histone H3 (H3Cit), and the nucleosome. The immunostaining revealed that DNA staining (DAPI), MPO, histone H3.1, histone H3Cit, and nucleosome signals colocalize in extruded NETs released from DMSO-dHL-60+PMA cells (Figure 1 A-D). Similarly, immunostaining on PMA-activated human neutrophils confirmed the colocalization of MPO, DNA, and H3.1 within NETs (Figure 1 E-F). Altogether, this colocalization indicates the close proximity of these components, suggesting they are part of the same structural entity.

**Figure 1.**
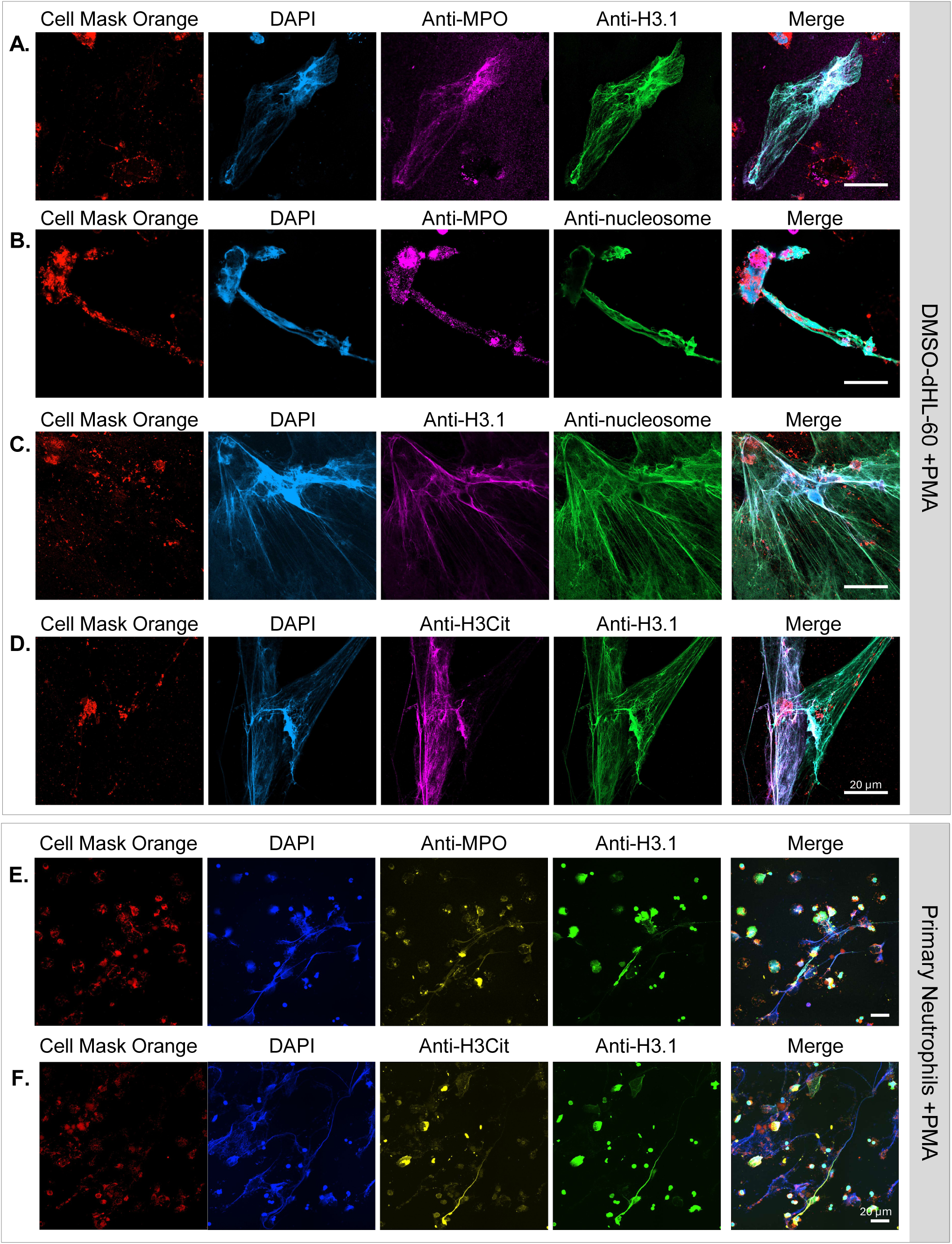
Immunostaining of NETs derived from HL-60 cells or primary neutrophils demonstrates the presence of H3.1-nucleosomes. Neutrophil-like DMSO-differentiated HL-60 cells (A-D) or primary neutrophils isolated from human whole blood (E-F) treated with PMA were stained with anti-MPO (A and B, purple; E, yellow), anti-histone H3.1 (A, D, E and F, green; C, purple), anti-nucleosome (B and C, green) or anti-H3Cit (D, purple; F, yellow) antibodies. DNA and membranes were stained with DAPI (blue) and Cell Mask^TM^ Orange (orange-red), respectively. Scale bars, 20µm. DMSO, dimethyl sulfoxide; PMA, phorbol 12-myristate 13-acetate; MPO, myeloperoxidase; NETs, neutrophil extracellular traps.

We further investigated, in this HL-60 NETs model, the interactions between MPO and H3.1-nucleosomes by immunoprecipitation (IP) followed by Western blots (Figure 2A). Characterization of the pulled-down proteins demonstrated that nucleosomes precipitated with either an anti-nucleosome antibody (IP anti-nucleosome) or an anti-H3.1 antibody (IP anti-H3.1) co-immunoprecipitated with the two subunits of MPO (MPOα and MPOβ), as well as histones H3 and H4. The different bands observed with the H3-antibody indicate the presence of various forms of histone H3, likely reflecting proteolytic cleavage events of this protein. Conversely, IP with anti-MPO-coated particles (IP anti-MPO) resulted in the co-IP of histones H3 and H4 as well as the two subunits of MPO (MPOα and MPOβ), suggesting that MPO and H3.1-nucleosomes interact closely in this NETs structure.

**Figure 2.**
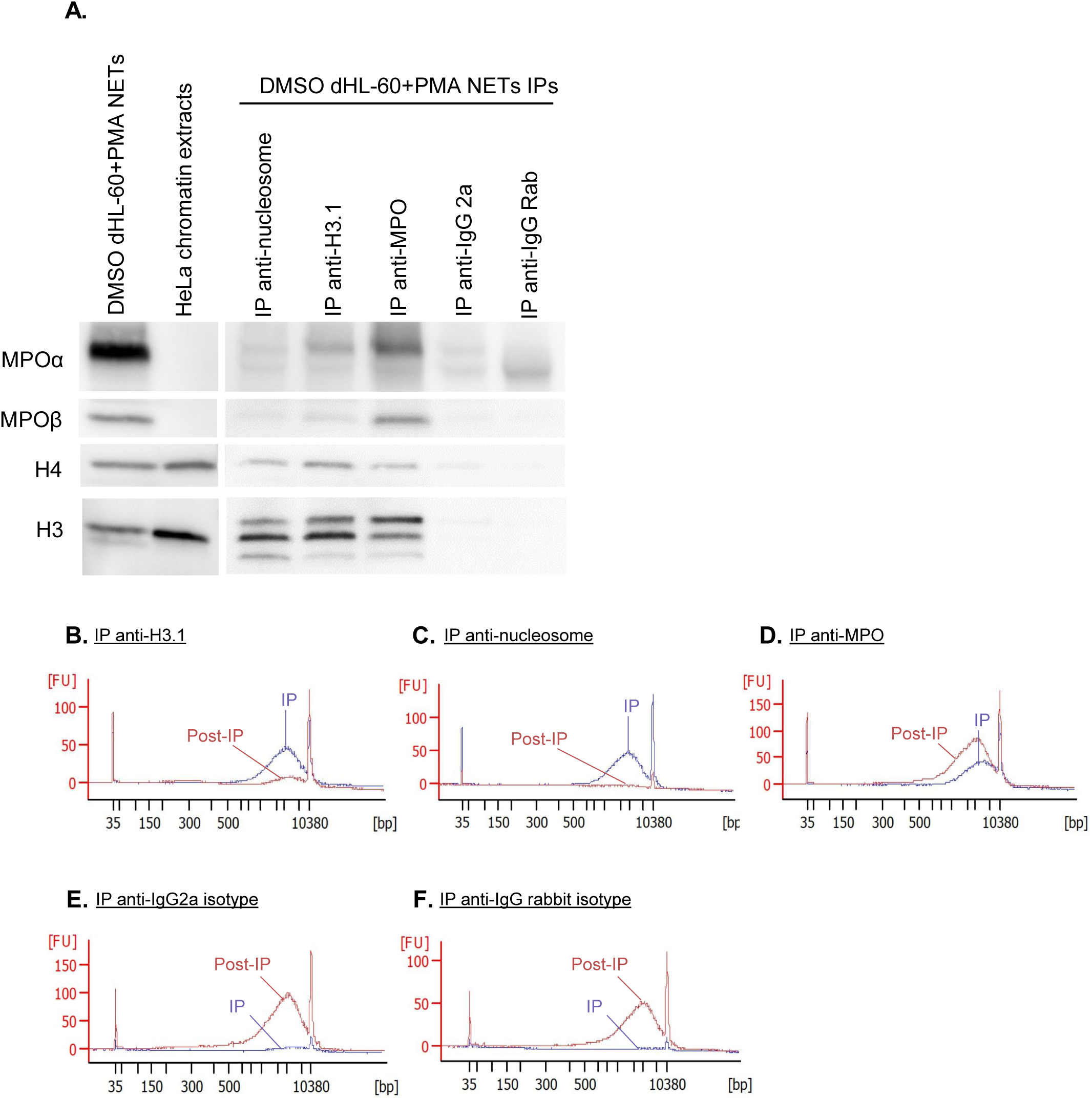
Molecular characterization of NETs from differentiated HL-60 cells demonstrates the presence of H3.1-nucleosomes. NETs extracts from DMSO-differentiated HL-60 cells treated with PMA (DMSO-dHL-60+PMA NETs) (A, lane 1), HeLa chromatin extracts (A, lane 2), and proteins pulled-down by using magnetic particles coated with anti-nucleosome (A, lane 3), anti-H3.1 (A, lane 4), anti-MPO (A, lane 5), anti-IgG2A (A, lane 6) and anti-IgG rabbit (A, lane 7) were analyzed using SDS-PAGE followed by anti-MPO, anti-H4 and anti-H3 Western blotting. The anti-MPO antibody detects both α and β sub-units of the MPO protein. The H3 antibody targets the C-terminal extremity of the histone H3. Cell-free DNA fragment size distribution of DNA immunoprecipitated (IP, blue) or left in the supernatant (post-IP, red) by anti-H3.1 (B), anti-nucleosome (C), anti-MPO (D), anti-IgG2A (E), and anti-IgG rabbit (F)-coated particles were analyzed using Agilent 2100 Bioanalyzer. DMSO, dimethyl sulfoxide; Ig, immunoglobulin; MPO, myeloperoxidase; NETs, neutrophil extracellular traps; PMA, phorbol 12-myristate 13-acetate; SDS-PAGE, sodium dodecyl-sulfate polyacrylamide gel electrophoresis.

Furthermore, to confirm that the H3.1-nucleosomes immunoprecipitated were part of NETs, we investigated their DNA fragment size profiles using Agilent 2100 Bioanalyzer. The analysis revealed that the DNA fragments immunoprecipitated with anti-H3.1-, anti-nucleosomes-, and anti-MPO-coated beads measured thousands of base pairs consistent with the large NETs filaments (Figure 2B-D). The DNA characterization of the material remaining in the supernatant after IP anti-H3.1 and IP anti-nucleosome (post-IP) showed an absence of DNA, confirming that these IPs effectively captured all NETs structures. In contrast, the IP anti-MPO demonstrated only partial retention of the DNA, indicating incomplete capture of the NETs structures in this condition. Control experiments performed with anti-immunoglobulin (Ig) G2A and anti-IgG rabbit isotypes magnetic particles did not significantly precipitate either MPO, histone H3, histone H4 (Figure 2A), or DNA (Figures 2E-F), confirming the specificity of the IPs.

Subsequently, to validate the interaction between the NETs components and H3.1-nucleosomes, we quantified the depletion of H3.1-nucleosomes following IP anti-nucleosome, IP anti-H3.1, and IP anti-MPO. Quantification results revealed a 97% depletion of H3.1-nucleosomes after IP anti-nucleosome, 99% depletion after IP anti-H3.1, and 30% depletion after IP anti-MPO, confirming the partial co-IP of H3.1-nucleosomes and MPO (S1 Table). IP with anti-IgG2a and anti-IgG isotype controls did not result in nucleosome depletion confirming the specificity of the IP.

Overall, our findings confirm interactions between H3.1-nucleosomes, MPO, and DNA, demonstrating the presence of H3.1-nucleosomes within NETs and highlighting the utility of H3.1-nucleosome quantification as a reliable marker for NETs detection.

### 3.2. Development of a ChLIA for quantification of H3.1-nucleosomes in plasma

We then developed and analytically validated an automated chemiluminescent magnetic bead-based immunoassay to measure H3.1-nucleosomes in K2EDTA plasma samples. This sandwich assay uses the anti-histone H3.1 antibody coated on magnetic particles and an anti-nucleosome conformational antibody in the liquid phase. The linear range for the assay was established between 26.3 ng/mL to 215.9 ng/mL (Mix 1, Figure 3A), 201.4 ng/mL to 971.8 ng/mL (Mix 2, Figure 3B), and 389.8 ng/mL to 1214.4 ng/mL (Mix 3, Figure 3C). Combining these results, the linear quantification range for the H3.1-nucleosome assay spans from 26.3 ng/ml to 1214.4 ng/mL (Figure 3D and S2 Table). This range was verified by fitting a linear model based on the three combined mixes (33 proportions; combined mixes, Figure 3D). Additionally, for samples with H3.1-nucleosome levels higher than 1200 ng/mL, we developed and validated an automatic 5-fold dilution to extend the range, resulting in a linear quantification range for the H3.1-nucleosome assay from 26.3 ng/mL to 6000 ng/mL. All limit of blank (LOB), limit of detection (LOD), and limit of quantification (LOQ) values estimated for three independent lots of reagents were below 3 ng/mL (mean ± standard deviation [SD]: LOB = 0.33 ± 0.06; LOD = 0.93 ± 0.06; LOQ = 2.30 ± 0.14), defined as the assay limit (Figure 3E).

**Figure 3.**
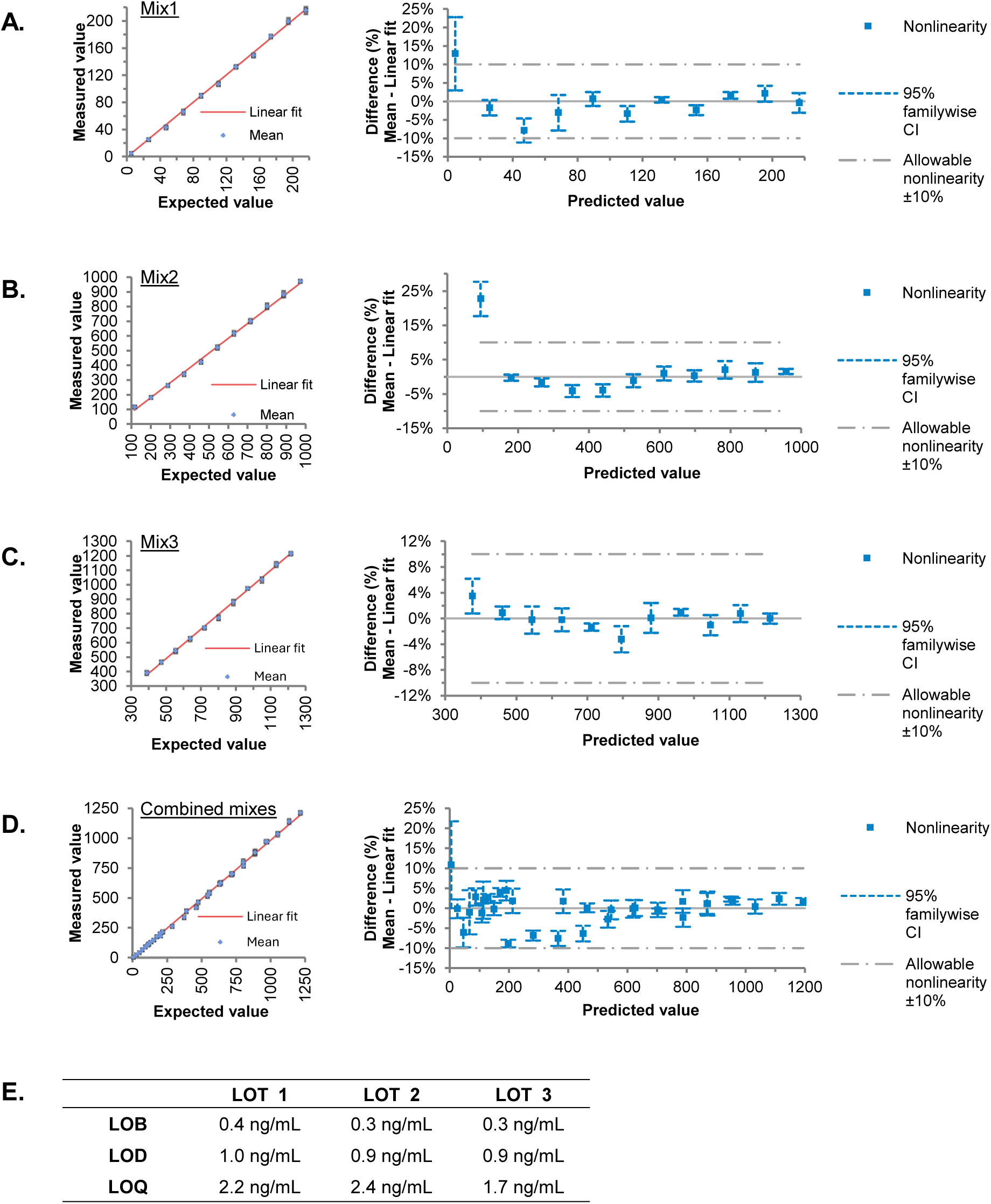
H3.1-nucleosome immunoassay linearity and sensitivity. Three K2EDTA plasma samples containing low concentrations of H3.1-nucleosomes were each mixed into three other K2EDTA plasma samples of higher H3.1 concentrations [(A) Mix 1, (B) Mix 2 or (C) Mix 3] to prepare 11 proportions and measured with the H3.1-nucleosomes immunoassay. Scatter plots show the linearity of the measurement, expressed in ng/mL, over the measuring intervals (expected value). The red line shows the linear fit. The linearity plots show the difference between the linear and nonlinear fit across the measuring interval. (D) Combined results from Mixes 1–3. (E) H3.1-nucleosomes assay limits of blank (LOB), limits of detection (LOB) and limits of quantification (LOQ) estimated for three independent lots of reagents. CI, confidence

Then, the assay precision was assessed, and quantification was performed either by calibrating the assay daily (Table 1A) or using a single calibration per reagent lot on the first day (Table 1B). Within-run coefficients of variation (CV) ranged from 1.4% to 2.8%, within-lot CV from 2.9% to 6.4%, and within-laboratory CV from 3.1% to 9.2%, depending on the sample considered. Interestingly, while bias in concentration between daily and single calibration was limited, single calibration induced slightly higher imprecision, though still reasonable.

**Table 1.**
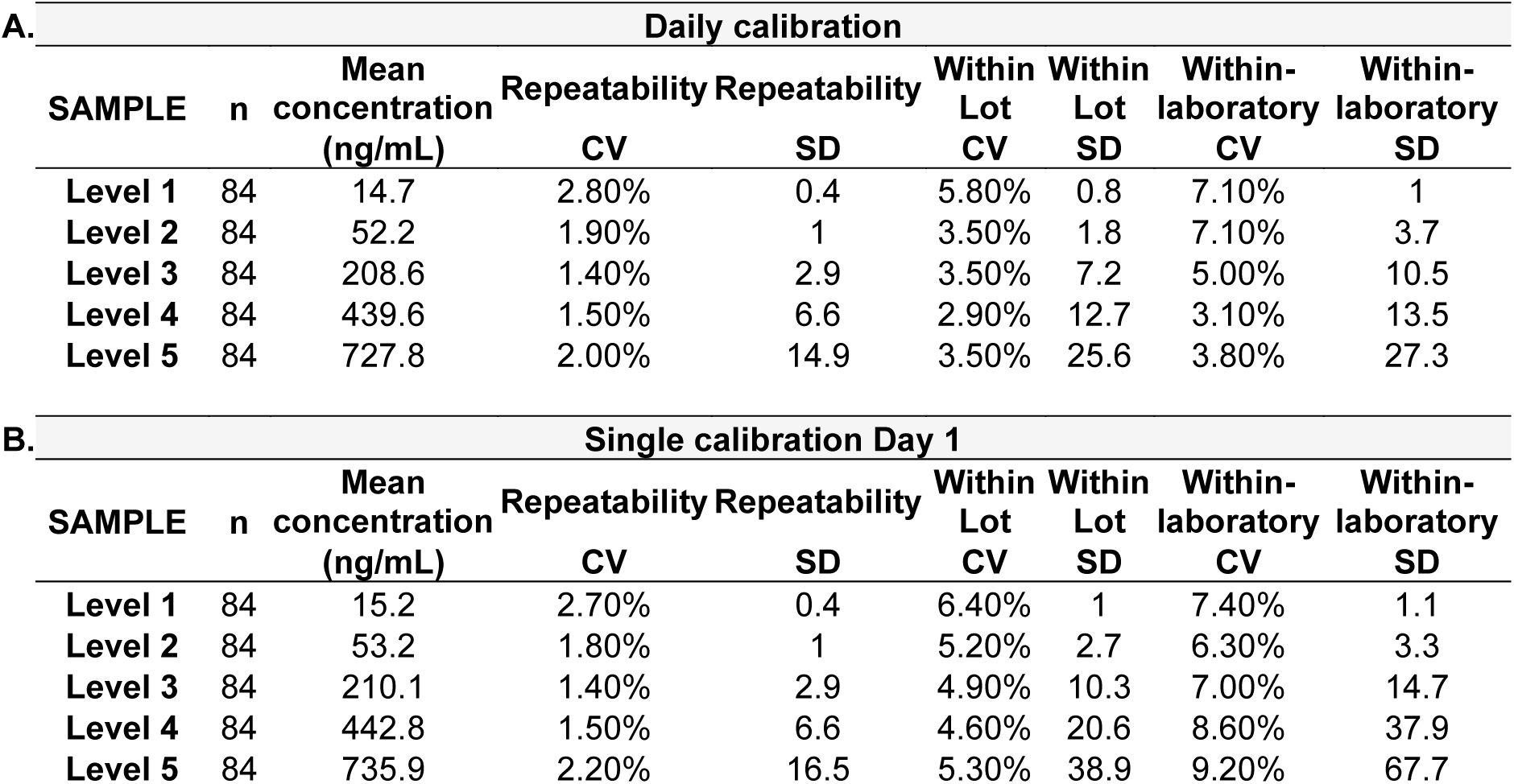
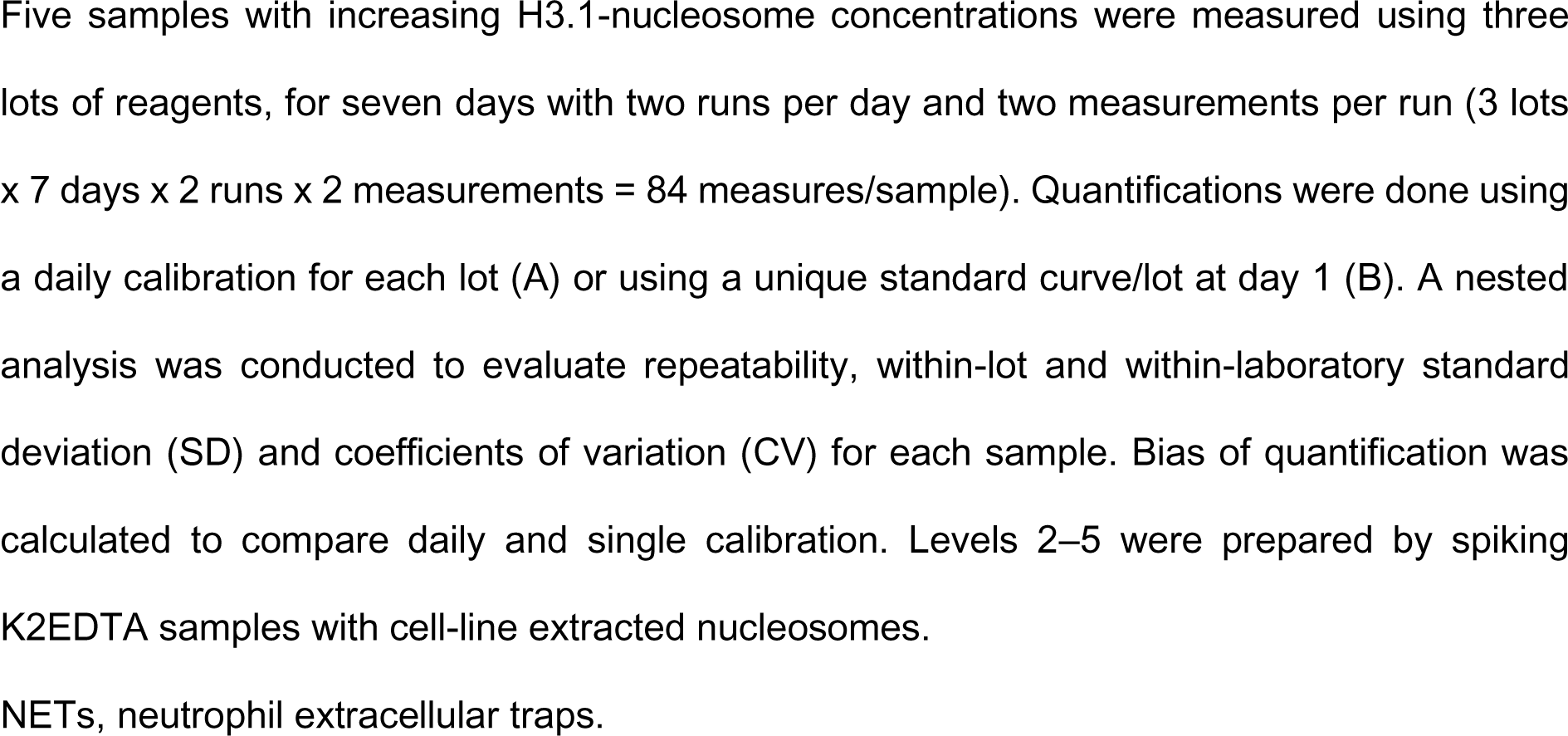
H3.1-nucleosome immunoassay precision.

Additionally, a spike-recovery study was performed by adding known concentrations of H3.1-nucleosome into three independent K2EDTA plasma samples. Recovery rates ranged from 89% to 105%, indicating no matrix effect (Table 2).

**Table 2.**
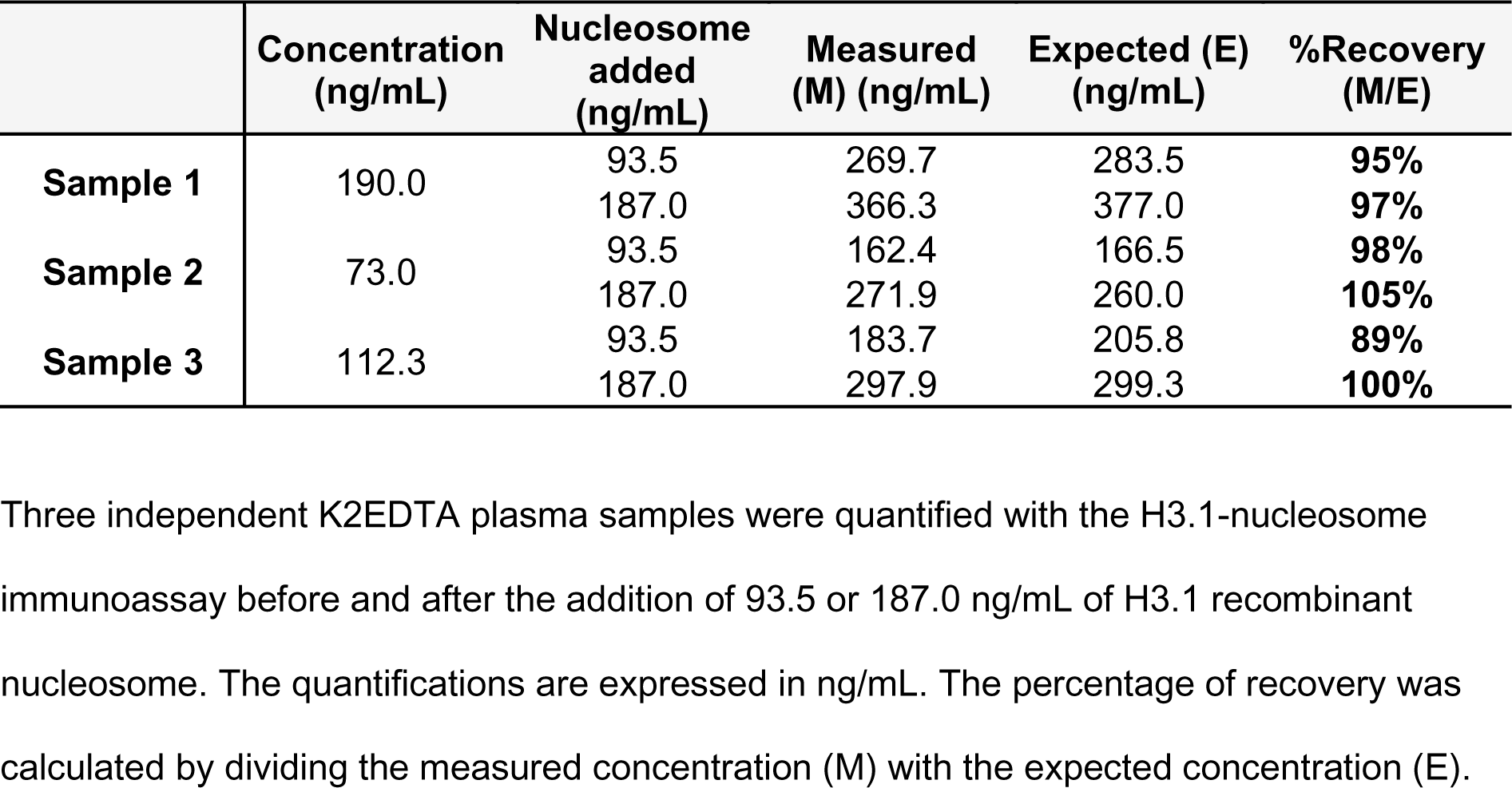
H3.1-nucleosome immunoassay spike recovery.

|  | Concentration<br>(ng/mL) | Nucleosome<br>added<br>(ng/mL) | Measured<br>(M) (ng/mL) | Expected (E)<br>(ng/mL) | %Recovery<br>(M/E) |
| --- | --- | --- | --- | --- | --- |
| **Sample 1** | 190.0 | 93.5 | 269.7 | 283.5 | **95%** |
|  |  | 187.0 | 366.3 | 377.0 | **97%** |
| **Sample 2** | 73.0 | 93.5 | 162.4 | 166.5 | **98%** |
|  |  | 187.0 | 271.9 | 260.0 | **105%** |
| **Sample 3** | 112.3 | 93.5 | 183.7 | 205.8 | **89%** |
|  |  | 187.0 | 297.9 | 299.3 | **100%** |

Three independent K2EDTA plasma samples were quantified with the H3.1-nucleosome immunoassay before and after the addition of 93.5 or 187.0 ng/mL of H3.1 recombinant nucleosome. The quantifications are expressed in ng/mL. The percentage of recovery was calculated by dividing the measured concentration (M) with the expected concentration (E).

Then, the specificity of the H3.1-nucleosome ChLIA assay for the H3.1-nucleosome structure was confirmed by testing the recombinant H3.3-nucleosome and the recombinant H3.1-histone, both exhibiting a cross-reactivity lower than 2%. Interestingly, when recombinant H3.1-nucleosomes presenting citrullinated marks (H3R8Cit and H3R2, 8,17Cit) were tested in the assay, the quantification yielded values of 88.5% and 113.8%, respectively, relative to H3.1-nucleosomes (S2 Figure). These results demonstrate that the presence of citrulline marks on histone H3 does not interfere with the accurate quantification of H3.1-nucleosomes. Afterwards, common endogenous interference substances from plasma were tested by comparing H3.1-nucleosome measurements with and without these substances in plasma samples. Table S3 shows concentrations of these substances that impact H3.1-nucleosome measurements less than 10%.

Overall, these analytical validation results confirm that the H3.1-nucleosome ChLIA assay is precise, sensitive, specific, and robust for measuring H3.1-nucleosome concentrations in K2EDTA plasma samples.

### 3.3. The H3.1-nucleosome assay effectively detects NETs-related nucleosomes

Prior to investigating the clinical relevance of the H3.1-nucleosome ChLIA assay, we assessed its efficacy in detecting NETs-related H3.1-nucleosomes in PMA-treated DMSO-differentiated HL-60 cells. The assay showed a significant increase of H3.1-nucleosome levels in DMSO-differentiated HL-60+PMA cells (here referred to as NETs; mean ± SD: 1306 ± 238 ng/mL) compared with the untreated DMSO-differentiated HL-60 cells (here referred to as controls; mean ± SD: 302 ± 175 ng/mL; ** p = 0.0054) (Figure 4A). In parallel, this assay’s performance was compared with other potential NETs biomarkers such as the extracellular double-stranded DNA (dsDNA), the MPO-DNA complex measured using a widely referenced custom ELISA assay (31) and nucleosomes containing citrullinated H3-histones (H3R8cit-nucleosome). While the H3R8cit-nucleosome assay differentiated between PMA-treated (mean ± SD: 454 ±138 ng/mL) and untreated DMSO differentiated HL-60 (mean ± SD: 134 ± 103 ng/mL), its significance was lower; (* p = 0.0364) than that of H3.1-nucleosomes (Figure 4B). Extracellular dsDNA concentrations were significantly higher in PMA-treated DMSO-differentiated HL-60 cells (mean ± SD: 10.15 ± 0.57 ng/µL) compared to untreated cells (mean ± SD: 2.97 ± 0.92 ng/µL; *** p = 0.0008) (Figure 4C). Although MPO-DNA complex signal (optical density [OD]) was elevated in PMA-treated DMSO-differentiated HL-60 extracts (mean OD value 1.42 ± 0.69) compared with the control conditions (mean OD value 0.51 ± 0.07; p = 0.1418), the difference was not statistically significant (Figure 4D). Our findings confirm that the H3.1-nucleosomes ChLIA assay effectively detects H3.1-nucleosomes associated with NETosis in *in-vitro* model.

**Figure 4.**
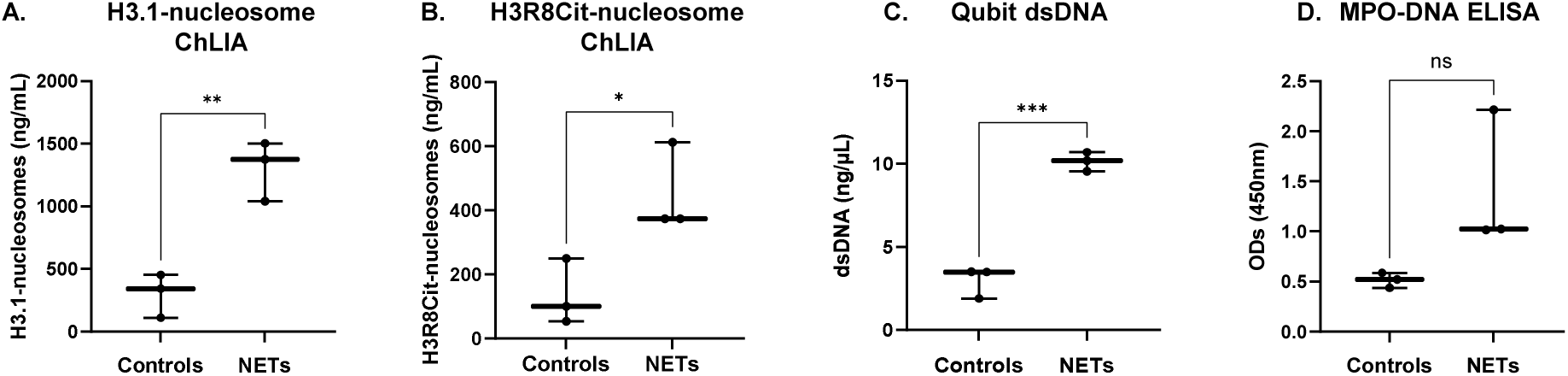
Comparative measures of H3.1-nucleosomes, H3R8Cit-nucleosomes, dsDNA and MPO-DNA complexes in NETs derived from HL-60 cells. Jitter plot analysis representation of the H3.1-nucleosomes (A), the H3R8Cit-nucleosomes (B), the dsDNA (C), and the MPO-DNA (D) measured for three independent productions of NETs extracts (n = 3) from DMSO-differentiated HL-60 cells treated with PMA (NETs) compared with the untreated cells (Controls). ns represent non-significant p-value, *, ** and *** represent p-values <0.05, <0.01 and <0.001, respectively, calculated using the Welch’s test. Cit, citrullinated histone H3; dsDNA, double-stranded DNA; MPO, myeloperoxidase; NETs, neutrophil extracellular traps; OD, optical density.

### 3.4. The H3.1-nucleosomes assay significantly distinguishes NETs-associated diseases from healthy controls

To demonstrate the clinical relevance of the H3.1-nucleosomes ChLIA assay, we conducted a study on a retrospective cohort (n = 538 patients) constituted by 302 K2EDTA plasma samples from patients presenting 11 different NETs-related pathologies and 236 K2EDTA plasma samples from healthy donors (n=236 donors). The results indicated significantly elevated levels of circulating H3.1-nucleosomes in patients with NETs-related pathologies (here referred to as NETs patients) compared with the control population (here referred to as Healthy donors - mean 721.00 ng/mL vs 30.66 ng/mL, **** p < 0.0001) (Figure 5A). Furthermore, to evaluate the performance of the assay in distinguishing patients suffering from NETs-related diseases vs heathy individuals, we generated a receiver-operating characteristic (ROC) curve. The area under the ROC curve (AUC) was calculated as 0.91 (95% CI: 0.89–0.94) (Figure 5B). A sensitivity of 85.8% at 86.9% of specificity was observed at the optimal threshold of 47.25 ng/mL, as defined using the Youden Index. The positive predictive value and the negative predictive value were 89.31% and 82.66%, respectively. The results of this study confirmed the clinical efficacy of the H3.1-nucleosomes ChLIA assay in detecting elevated levels of circulating nucleosomes in patients with NETs-related pathologies. The high sensitivity and specificity, combined with robust predictive values, highlight the potential of this assay for clinical use in identifying NETs-related conditions.

**Figure 5.**
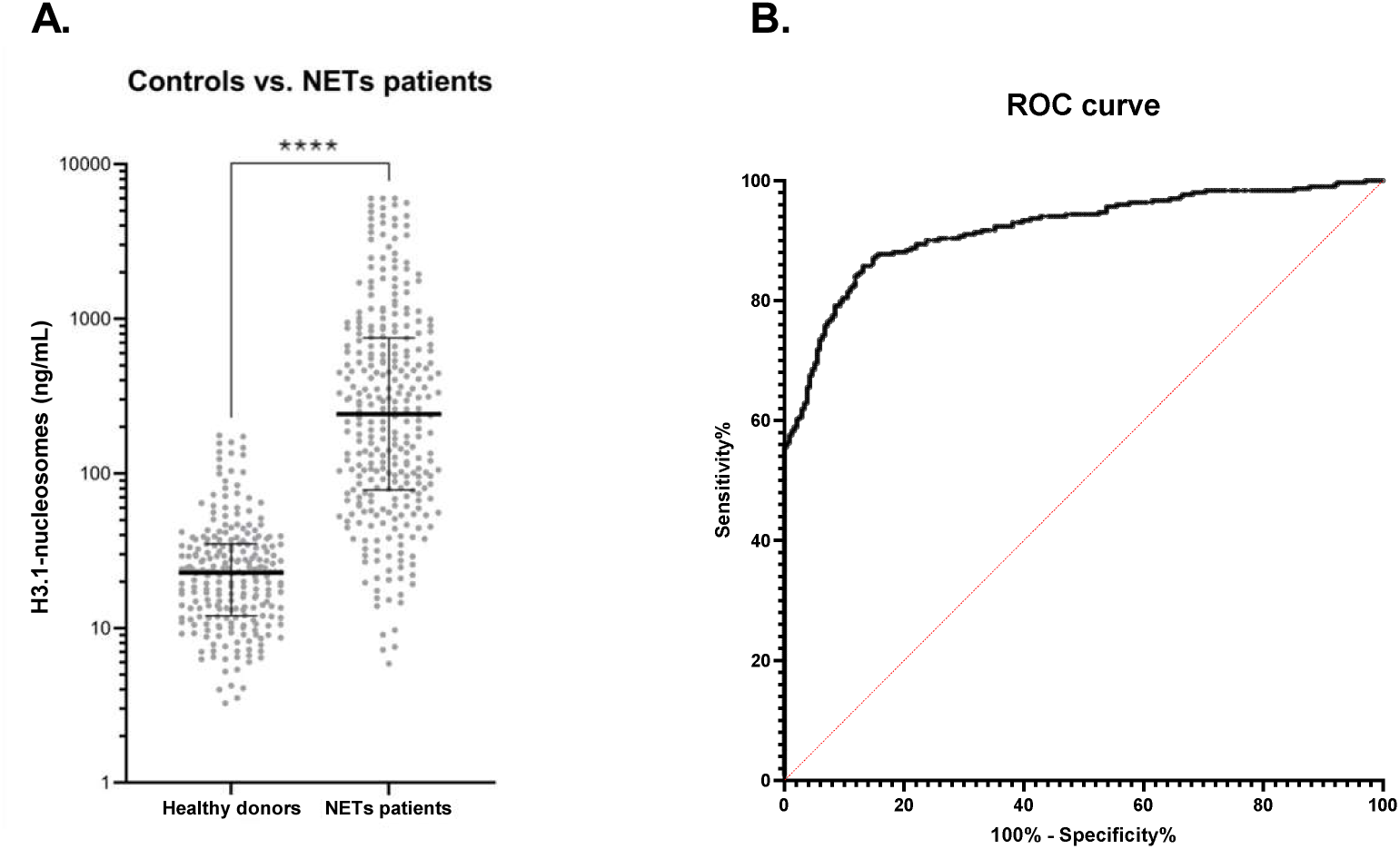
The H3.1-nucleosome assay significantly distinguishes NETs-associated diseases from healthy controls. (A) Jitter plot analysis representation of the circulating H3.1-nucleosomes level (ng/mL). The whiskers represent the 25th–75th percentile with median. **** represent p-value <0.0001, calculated using the Mann–Whitney U test. (B) ROC curve analysis of circulating H3.1-nucleosomes for discrimination of NETs patient’s vs control samples. The area under the curve is 0.91 (95% CI: 0.89–0.94; p < 0.0001). CI, confidence interval; NETs, neutrophil extracellular traps; ROC, receiver operating characteristic.

## 4. Discussion

During NETosis, neutrophils release their decondensed DNA associated with histones and host defense proteins such as MPO and NE into the extracellular space to trap pathogens (9, 45). However, excessive release of NETs or dysfunction in their clearance can cause detrimental effects and have been described in various pathologies including sepsis, COVID-19, rheumatoid arthritis, systemic lupus erythematosus, and others. This highlights the clinical relevance of monitoring NETs in blood samples. Given the central role of NETs in these diseases, their quantification could serve as a biomarker and/or risk factor for disease severity, potentially aiding personalized treatment. Based on this clinical evidence, the process of NET formation has attracted increasing attention from researchers, but the absence of standardized NET quantification method complicates cross-laboratory comparisons and highlights the need for consistent methodologies.

Current techniques for NET quantification primarily rely on immunostaining (46-50); but this method presents important limitations for clinical studies, especially in liquid biopsy (51). For instance, the major limitation of the immunostaining is its time-consuming nature, and prone to manual counting bias, which can lead to variability among individuals evaluating the same images. Although this limitation has been mitigated by automated NET-based fluorescence microscopy assays (52), which have minimized observer subjectivity, these methods remain laborious and susceptible to residual bias. To overcome these limitations, ELISA-based approaches are also widely used to detect NETs (29, 32, 45, 53, 54) mainly by targeting NETs components such as MPO, NE, or histones (e.g. H3Cit) using specific capture antibodies paired with an anti-DNA detection antibody. Despite the relevance of the results obtained using these immunoassays, those are often not fully quantitative, with results presented as absorbance values or relative fluorescence units (32), or quantified using an “in-house” standard curve (30), making cross-laboratory comparison difficult.

In this study, we investigated the presence of H3.1-nucleosomes in NETs. To this end, we used the *in vitro* HL-60 model (41, 42) and primary human neutrophils (43), both treated with PMA to induce NETosis, and we confirmed the presence of H3.1-nucleosomes in the resulting NETs. Immunostaining experiments revealed colocalization of histones H3.1 and H3Cit, DNA, and MPO, suggesting a close interaction among these components as well with nucleosomes, indicating that H3.1-nucleosomes are part of the NETs structure. Molecular characterization further corroborated these findings, with co-IP experiments identifying complexes of MPO, histones H3 and H4, and DNA. Notably, substantial DNA levels were detected in the post-capture supernatant of anti-MPO IP experiments, potentially indicating incomplete IP or the presence of MPO unbound to DNA, as supported by other studies (55-58). Given this, along with the fact that MPO is required for PMA-induced but not bacterial-induced NETs formation (58), and that MPO can be released into the extracellular space during degranulation (59), MPO-DNA quantification may not be the most suitable biomarker for clinical applications. Additionally, it is important to acknowledge that PMA induces a specific subset of NETs, which may not fully represent all forms of NETs observed in physiological conditions. Despite this limitation, our *in vitro* approaches successfully demonstrated that H3.1-nucleosomes are associated with NETs structure.

To specifically quantify H3.1-nucleosomes, we developed a chemiluminescent immunoassay, adhering to Clinical & Laboratory Standards Institute (CLSI) guidelines. This analytically validated, automated assay enables high-sensitivity quantification of circulating H3.1-nucleosomes using only 50 µl of K2EDTA plasma samples and includes an internal calibrant to ensure consistency and comparability across different laboratories.

If it appears clearly that H3.1-nucleosomes are key chromatin components and major constituents of NETs, histone citrullination mediated by peptidyl arginine deiminase is another widely studied NET biomarker (60-64). Although citrullinated histones are recognized as key NET components (65, 66), not all NETosis pathways involve citrullination, limiting their reliability as NET biomarkers (67, 68). In contrast, the H3.1-nucleosome assay is capable of detecting H3.1-nucleosomes regardless of post-translational modifications on histone H3 tails, including methylation, acetylation, phosphorylation, or citrullination (69). This makes it a potentially more universal tool for NET detection. Moreover, neutrophil protease activity during NETosis may cleave histone H3, removing citrullinated marks from the N-terminal tail (70). Consistently, our Western Blot characterization using an anti-H3 C-terminal antibody detected multiple bands in PMA-treated DMSO-differentiated HL-60 cells, indicating N-terminal cleavages of histone H3. However, it is essential to admit that circulating H3.1-nucleosomes are not exclusive to NETosis and may arise from other cell death forms, such as apoptosis or necrosis, or from other cell types besides neutrophils. Nonetheless, the pronounced increase of H3.1-nucleosomes observed in NET-related diseases, compared to healthy controls or patients suffering from non-NETs related diseases (e.g., cancers without significant systemic inflammation) underscores the relevance of the H3.1-nucleosome assay as an indicator of NETosis (71).

Based on this evidence, we evaluated the detection of H3.1-nucleosome by an immunoassay as a potential breakthrough method for objective, robust, reproductible and quantitative NETs detection. In PMA-treated DMSO-differentiated HL-60 cells, the assay detected elevated H3.1-nucleosome levels, correlating with established NET markers, such as MPO-DNA, H3R8cit-nucleosome and extracellular dsDNA. This consistency was also observed in whole blood NETosis models (72), further validating the H3.1-nucleosome assay as a reliable indicator of NETosis.

Finally, by measuring H3.1-nucleosome levels in clinical K2EDTA plasma samples from both healthy donors and patients suffering from NETs-related diseases, we detected basal levels of H3.1-nucleosomes in healthy donors, reflecting the basal nucleosome release from dying and stressed cells into the blood circulation (4), while significantly elevated levels were detected in patients with NET-related diseases, demonstrating the potential clinical utility of the H3.1-nucleosomes assay as a robust tool for NETs detection. Further clinical studies will be required to validate H3.1-nucleosome measurements for risk prediction and prognosis in NET-related diseases such as sepsis.

## Data Availability statement

The raw data supporting the conclusions of this article will be made available by the authors, without undue reservation.

## Author contributions

Conceptualization, M.H., M.W., P.V.d.A.; Methodology, M.H., M.W., P.V.d.A., G.R., D.P., J.C. F.L.; Validation, G.R., F.J., J.C., D.P., R.V.; Formal Analysis, G.R., D.P., J.C.; Investigation, M.W., P.V.d.A., G.R., R.V., A.L., F.J., O.T., F.L., V.L., F.S.; Writing – Original Draft, M.W., P.V.d.A., G.R., J.C., D.P. Writing – Review & Editing; M.W., G.R., J.C., D.P., R.V., A.L., F.J., V.L., O.T., F.L., F.S., P.V.d.A., M.H.; Visualization, G.R., M.W., A.L., J.C.; Supervision, M.H.; Project administration, M.W., G.R., M.H. All authors approved the final manuscript.

## Acknowledgements

We acknowledge Marie Lurkin for her assistance in the processing of samples using the immunoassay automated platform and the Logistics team of Belgian Volition for their support. We thank Terry Kelly for her careful proofreading of the manuscript. We also thank the MorphIM Platform at the University of Namur for their expert guidance and assistance in generating and analyzing fluorescent microscopy images. Microscopy and image analysis of primary neutrophils was performed at the Nikon Imaging Center at UC San Diego. We’d like to thank Richard Sanchez and the Nikon Imaging Center at UCSD for the support on microscopy experiments.

## Supporting information captions

**S1 Figure: PMA treatment induces NET formation in DMSO-differentiated HL-60 cells**

Neutrophil-like DMSO-differentiated HL-60 cells were treated during 5 hours with the NETosis-inducer PMA (PMA, lower panel), or not treated (DMSO, upper panel). DNA and membranes were stained with DAPI (Blue) and Cell Mask Orange (Orange-Red), respectively. Scale bars 50 µm.

**S2 Figure: Detection of citrullinated H3.1-nucleosomes using the H3.1-nucleosome immunoassay**

Quantification of H3.1-, H3R8Cit-, and H3R2,8,17Cit-recombinant nucleosomes using the H3.1-nucleosome immunoassay. Results are expressed as a percentage relative to the theoretical concentration of nucleosomes loaded in the samples (% recovery).

**S1 Table: H3.1-nucleosomes depletion after immunoprecipitation**

H3.1-nucleosomes levels, expressed in ng/mL, indicate the depletion of nucleosomes after immunoprecipitation (IP) in comparison to the level present in the initial samples (HL-60 NETs unprocessed). Depletion results are expressed in %.

NETs, neutrophil extracellular traps.

**S2 Table: H3.1-nucleosome immunoassay linearity**

NETs, neutrophil extracellular traps; Mean cc, mean of H3.1-nucleosomes assay concentration expressed in ng/mL.

**S3 Table: H3.1-nucleosome immunoassay interferences and cross-reactivity**

NETs, neutrophil extracellular traps

**S4 Table: Descriptive statistics**

NETs, neutrophil extracellular traps; ROC, receiver operating characteristic.

**S1 Raw Images: Western blots raw images used for Figure 2A**

