## Supporting information for "Quantification of H3.1-nucleosomes using a chemiluminescent immunoassay: a reliable method for neutrophil extracellular trap detection"

### S1 Figure: PMA treatment induces NETs formation in DMSO-differentiated HL-60 cells

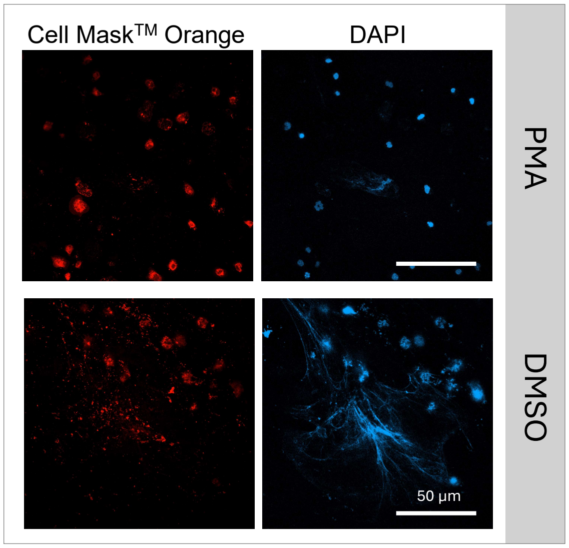


**Supplementary Figure 1 :** Neutrophil-like DMSO-differentiated HL-60 cells were treated during 5 hours with the NETosis-inducer PMA (PMA, lower panel), or not treated (DMSO, upper panel). DNA and membranes were stained with DAPI (Blue) and Cell Mask Orange (Orange-Red), respectively. Scale bars 50 µm.

S2 Figure: Detection of citrullinated H3.1-nucleosomes using the H3.1-nucleosome immunoassay

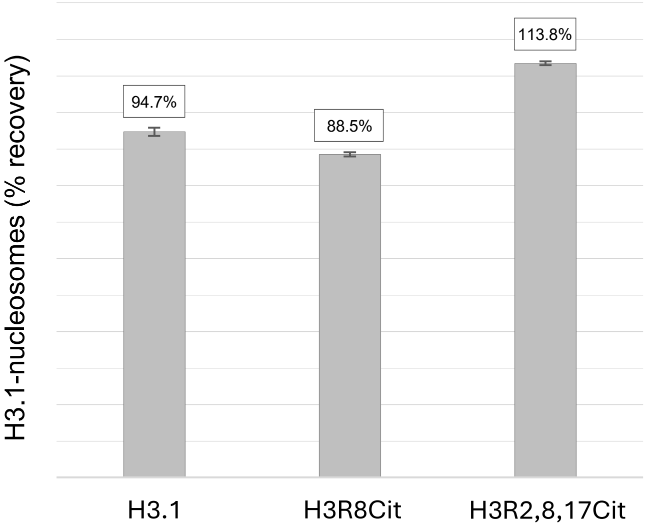


**Supplementary Figure 2:** Quantification of H3.1-, H3R8Cit-, and H3R2,8,17Cit-recombinant nucleosomes using the H3.1-nucleosome immunoassay. Results are expressed as a percentage relative to the theoretical concentration of nucleosomes loaded in the samples (% recovery).

### S1 Table: H3.1-nucleosomes depletion after immunoprecipitation

| Samples | H3.1-nucleosome concentration post IP (ng/mL) | % depletion |
|---|---|---|
| HL-60 NETs unprocessed | 468.4 | - |
| IP anti-nucleosome | 12.5 | 97% |
| IP anti-H3.1 | 5.7 | 99% |
| IP anti-MPO | 328.0 | 30% |
| ÏP anti-IgG2a isotype | 458.1 | 2% |
| IP anti-IgG rabbit isotype | 490.7 | -5% |

**Supplementary Table 1:** H3.1-nucleosome levels, expressed in ng/mL, showing the depletion of H3.1-nucleosomes after immunoprecipitation in comparison to the level present in the initial samples (dHL-60 NETs unprocessed). Depletion results are expressed in %.

S2 Table: H3.1-nucleosome immunoassay linearity

| | Mix 1 | | | | Mix 2 | | | | Mix 3 | | | |
|---|---|---|---|---|---|---|---|---|---|---|---|---|
| | Expected value | Mean cc | Linear fit | Non linearity | Expected value | Mean cc | Linear fit | Non linearity | Expected value | Mean cc | Linear fit | Non linearity |
| **Sample 1 (S1)** | **5.2** | 5.2 | 4.6 | **12.9%** | **115.8** | 115.8 | 94.4 | **22.7%** | **389.8** | 389.8 | 376.7 | **3.5%** |
| **90% S1 - 10% S2** | 26.3 | 25.4 | 25.8 | **-1.8%** | 201.4 | 180.2 | 180.7 | **-0.3%** | 472.2 | 464.5 | 460.5 | **0.9%** |
| **80% S1 - 20% S2** | 47.3 | 43.3 | 47.0 | **-7.9%** | 287.0 | 262.6 | 266.9 | **-1.6%** | 554.7 | 543.0 | 544.3 | **-0.2%** |
| **70% S1 - 30% S2** | 68.4 | 66.2 | 68.3 | **-3.1%** | 372.6 | 338.5 | 353.2 | **-4.2%** | 637.2 | 626.7 | 628.1 | **-0.2%** |
| **60% S1 - 40% S2** | 89.5 | 90.0 | 89.5 | **0.6%** | 458.2 | 421.8 | 439.5 | **-4.0%** | 719.6 | 702.3 | 711.9 | **-1.3%** |
| **50% S1 - 50% S2** | 110.5 | 107.0 | 110.7 | **-3.4%** | 543.8 | 519.6 | 525.8 | **-1.2%** | 802.1 | 770.0 | 795.7 | **-3.2%** |
| **40% S1 - 60% S2** | 131.6 | 132.5 | 131.9 | **0.4%** | 629.4 | 617.8 | 612.1 | **0.9%** | 884.5 | 880.0 | 879.5 | **0.1%** |
| **30% S1 - 70% S2** | 152.7 | 149.5 | 153.2 | **-2.4%** | 715.0 | 700.1 | 698.4 | **0.2%** | 967.0 | 972.5 | 963.3 | **1.0%** |
| **20% S1 - 80% S2** | 173.7 | 177.2 | 174.4 | **1.6%** | 800.6 | 800.3 | 784.7 | **2.0%** | 1049.4 | 1036.0 | 1047.1 | **-1.1%** |
| **10% S1 - 90% S2** | 194.8 | 199.7 | 195.6 | **2.1%** | 886.2 | 881.5 | 871.0 | **1.2%** | 1131.9 | 1139.3 | 1130.8 | **0.7%** |
| **Sample 2 (S2)** | **215.9** | 215.9 | 216.8 | **-0.5%** | **971.8** | 971.8 | 957.3 | **1.5%** | **1214.4** | 1214.4 | 1214.6 | **0.0%** |

S3 Table: H3.1-nucleosome immunoassay interferences and cross-reactivity

| Potentially interfering agent | Threshold concentration |
|---|---|
| Haemoglobin | 50 mg/dL |
| Bilirubin conjugate | 20 mg/dL |
| Bilirubin non conjugated | 20 mg/dL |
| Protein | 8 g/dL |
| Triglyceride (intralipid) | 3000 mg/dL |
| Cholesterol | 300 mg/dL |
| HAMA (Human anti-mouse antibodies) | 600 ng/mL |
| Rheumatoid Factor | 2940 IU/mL |

S4 Table: Descriptive statistics

| | Control donors | « NETs » patients |
|---|---|---|
| **Number of samples** | 236 | 302 |
| **Descriptive statistics (Nu.Q® H3.1 – ng/mL)** | | |
| Minimum | 3.26 | 5.89 |
| 25% Percentile | 12.02 | 78.07 |
| Median | 22.81 | 242.1 |
| Mean | 30.66 | 721 |
| 75% Percentile | 35 | 751.3 |
| Maximum | 175.9 | 6000 |
| Range | 172.7 | 5994 |
| **95% Confidence interval of median** | | |
| Actual confidence level | 95.66% | 95.62% |
| Lower confidence limit | 19.57 | 173.6 |
| Upper confidence limit | 24.45 | 308.8 |
| **Area under the ROC curve** | 0.9193 | |
| Std. Error | 0.01174 | |
| 95% confidence interval | 0.8963 to 0.9423 | |
| P-value | <0.0001 | |
| **Threshold at 65 ng/mL** | | |
| Sensitivity % | 79.14% | |
| Specificity % | 91.53% | |
